# CRISPR/Cas13a signal amplification linked immunosorbent assay (CLISA)

**DOI:** 10.1101/781237

**Authors:** Qian Chen, Tian Tian, Erhu Xiong, Po Wang, Xiaoming Zhou

**Author notes:** The authors declare no competing financial interest.

## Abstract

The enzyme-linked immunosorbent assay (ELISA) is a basic technique used in analytical and clinical investigations. However, conventional ELISA is still not sensitive enough to detect ultra-low concentrations of biomarkers for the early diagnosis of cancer, cardiovascular risk, neurological disorders, and infectious diseases. Herein we show a mechanism utilizing the CRISPR/Cas13a-based signal export amplification strategy, which double-amplifies the output signal by T7 RNA polymerase transcription and CRISPR/Cas13a collateral cleavage activity. This process is termed the CRISPR/Cas13a signal amplification linked immunosorbent assay (CLISA). The proposed method was validated by detecting an inflammatory factor, human interleukin-6 (human IL-6), and a tumor marker, human vascular endothelial growth factor (human VEGF), which achieved limit of detection (LOD) values of 45.81 fg/mL (2.29 fM) and 32.27 fg/m (0.81 fM), respectively, demonstrating that CLISA is at least 10^2^-fold more sensitive than conventional ELISA.

## INTRODUCTION

Immunoassays can be utilized for detecting almost any biomolecules, including proteins, small molecules, vesicles, nucleic acids, and even whole cells^1–2^. Since the invention of this method in the 1960s, immunoassays have undergone a developmental phase from radioimmunoassays to enzyme-linked immunoassays^3^. Using enzymes rather than radioactivity as the reporter label, enzyme-linked immunoassays, also termed enzyme-linked immunosorbent assays (ELISAs), have become the most widely used technique in both fundamental and applied immunological research^3^. With enzymatic signal amplification, ELISA has achieved a limit of detection (LOD) of approximately 0.01–50 ng/mL (pM to nM), depending on the affinity of the antibody^4^. Although it has achieved wide adoption in conventional diagnostic applications, ELISA is still not sensitive enough to detect ultra-low concentrations of biomarkers in the early diagnosis of cancer, cardiovascular risk, neurological disorders, and infectious diseases^5^.

To enhance the sensitivity of ELISA, current research has focused on improving the activity of enzymes, such as the currently popular nanozymes^6–7^. In addition, the researchers have explored nanoprobes to load enzymes due to their large specific surface area, which can increase the load of enzyme and achieve signal amplification^8–9^. However, since the nanomaterial is a non-biological material, it may impair the enzyme activity due to the low biocompatibility. Further, the nonuniformity of the nanoparticle may lead to a great error in measurements. Instead of an enzyme, a reporter using a DNA sequence as a signal output can significantly improve the sensitivity of the ELISA method. Typical examples include immuno-PCR^10–12^, immuno-RCA^13–14^, immuno-HCR^15–16^, proximity ligation assays^17–18^, and T7 transcription amplification^19^. As a consequence, the LOD of a given ELISA is, in general, enhanced 10~10^4^-fold by the use of DNA as a signal amplification element. Even with these developments, there are still significant challenges for their widespread adoption in analytical and clinical investigations. Possible limiting factors include the inability to achieve quantitative detection due to nonlinear signal amplification, which requires additional testing equipment and detection steps and, thus, is incompatible with existing commercial ELISA platforms.

CRISPR/Cas13a has been recently demonstrated to have RNA-directed RNA cleavage ability^20–22^. This RNA-guided trans-endonuclease activity is highly specific, being activated only when the target RNA has the perfect complementary sequence to the crRNA, and highly efficient (at least 10^4^ turnovers per target RNA recognition)^20–22^. This potent signal amplification ability of CRISPR/Cas13a enables the development of direct RNA assays with a sensitivity down to the fM level^20, 22–23^. Single molecule RNA detection could also be achieved when combined with a digital droplet assay^24^. Although there has been extensive development in nucleic acid detection, a CRISPR/Cas13a system has not yet been explored as an exciting opportunity for an immunoassay.

Herein, we report a new version of ELISA performed via the utilization of CRISPR/Cas13a as a signal export amplification strategy, which double-amplifies the output signal by T7 RNA polymerase transcription and CRISPR/Cas13a collateral cleavage activity: this strategy is called the CRISPR/Cas13a signal amplification linked immunosorbent assay (CLISA). It is the first example, to our knowledge, of the construction of a highly sensitive immunoassay based on a CRISPR technique.

## EXPERIMENTAL SECTION

### Experimental materials and instruments

All chemicals were purchased from the Guangzhou Chemical Reagent Factory. All DNA sequences were synthesized by Sangon Biotech (Shanghai), and fluorescent double-labeled probes were synthesized by Takara (Japan). All of the DNA and RNA sequences are provided in Table S1. T7 RNA polymerase, NTP, and transcription buffer were from Bio-Lifesci (Guangzhou, China). Human IL-6 and VEGF antigens and antibodies were purchased from KEY-BIO (Beijing, China). Commercial human IL-6 and human VEGF ELISA kits were purchased from ExCell Biotech (Shanghai, China). The reagents for protein expression and purification were obtained from Abiotech (Jinan, China). The plasmid used to express LbuCas13a is a gift from Professor Wang Yanli (Institute of Biophysics, Chinese Academy of Sciences). Fluorescence detection was performed on a SpectraMax iD5 multi-mode microplate reader (Molecular Devices). The PAGE electrophoresis experiments were performed using an instrument (Beijing Liuyi). All other solutions and buffers were prepared using ultrapure water (> 18.25 MΩ).

### Expression and purification of LbuCas13a proteins

*Escherichia coli Rosetta 2* (DE3) cells were cultured overnight in Terrific Broth containing chloramphenicol and ampicillin to express the LbuCas13a protein. Then, IPTG was added and induced at 37 °C for 4 h. The collected bacterial solution was centrifuged to obtain a precipitate and lysed by sonication in 20 mM Tris-HCl, pH 7.5, containing 1 M NaCl, 20 mM imidazole, and 10% glycerol. The supernatant was purified using a nickel column (Abiotech, Jinan, China) and eluted with eluent 1 (20 mM Tris-HCl, pH 7.5, 250 mM imidazole, and 150 mM NaCl) after centrifugation at 4 °C Then the eluted LbuCas13a protein was purified again using a heparin column (Abiotech, Jinan, China), and the protein was eluted with eluent 2 (20 mM Tris-HCl, pH 7.5, 1 M NaCl, and 10% glycerol). Finally, the protein was collected, and glycerol was added to a final concentration of 50%. The protein was stored at −80 °C for further use (Figure S1B).

### In Vitro Transcription of crRNA

The crRNA was prepared by in vitro transcription. The crDNA1 and crDNA2 templates were heated to 95 °C and then slowly cooled to room temperature for annealing. In a 50μL transcription system, 250 U T7 RNA polymerase, NTPs (final concentration of 0.5 mM), 50 Uμ recombination RNase inhibitor and 200 ng of template DNA were added and incubated for 4 h at 37 °C. Then, DNase I was added to digest the excess DNA template. Finally, the product of this reaction was purified by an RNA purification kit and stored at −20 °C for further use (Figure S1A).

### CLISA reaction

Dilutions of the specific antibody protein (5 μg / mL) in a coating buffer (carbonate-bicarbonate buffer, pH 9.6) were added to the plate, 0.1 mL per well, and incubated at 4 °C overnight. Then, the plate was blocked by adding 1% BSA protein, 0.3 mL per well, and incubated at 37 °C for 1 h. Serial dilutions of antigen were added to the plate, 0.1 mL per well, and incubated at 37 °C for 1 h.

Diluted biotinylated detection antibody (50 ng / mL) was added to the plate, 0.1 mL per well, and incubated at 37 °C for 1 h. Streptavidin and a biotinylated dsDNA amplification template were added to the plate one by one (0.1 mL per well) and incubated at 37 °C for 0.5 h. The non-bound solution was removed, and the wells were washed five times with PBS buffer containing 0.05% Tween-20 between each binding incubation. Finally, 50 μL of the reaction mixture (1×T7 buffer, 150 U T7 RNA polymerase, 1.25 mM NTPs) was added to each well and reacted at 37 °C for 1 h. Then, 100 μL of the Cas13a reaction system (final concentration of 100 nM LbuCas13a, 200 nM crRNA, 200 nM RNA fluorescent probe) was performed in different concentrations of the antigen wells. Fluorescent signals were recorded on a SpectraMax iD5 multi-function microplate reader. The reaction was conducted at 37 °C for 30 min and the fluorescence signal was recorded every minute.

For simplified CLISA detection, streptavidin was directly coated on 96-well plates. Then, the plate was blocked with 1% BSA protein, and a biotinylated dsDNA amplification template was added to the plate (0.1 mL per well) and incubated at 37 °C for 0.5 h. Next, we carried out the standard CLISA method.

### ELISA reaction

Serial dilutions of antigen were added to the plates at an amount of 100 μL per well following the protocol of the commercial kit and incubated at 37 °C for 1.5 h. Diluted biotinylated antibody L/well) was added to each well and incubated at 37 °C for 1 h. Next, diluted enzyme-binding working solution (100 μL /well) was added to each well and incubated at 37 °C for 30 minutes in the dark. The plate was washed five times with wash solution between each binding incubation. Finally, a chromogenic substrate (100 μL /well) was added and incubated at 37 °C for 15 min in the dark. Stop solution (100 μL /well) was added and the OD450 value (within 10 min) was measured immediately after mixing.

## RESULTS AND DISCUSSION

Prior to carrying out the CLISA, the collateral cleavage activity of CRISPR/Cas13a and the sensitivity of detecting the template RNA were first demonstrated. The CRISPR/Cas13a cleavage mechanism is shown in Figure 1A. Cas13a exhibits high activity under the guidance of crRNA in the presence of a synthesized target RNA. As shown in Figure 1B, the collateral cleavage activity of Cas13a can only be activated when Cas13a/crRNA/target RNA are present simultaneously (red curve). After that, we performed the CRISPR/Cas13a assay for RNA detection. As shown in Figure 1C, with the increase of target RNA concentrations, the fluorescence signals enhanced gradually, and the CRISPR/Cas13a system was able to detect the target RNA at as low of a concentration of 50 fM. Figure 1D shows an enlarged view of the low concentration measurement curve in Figure 1C. Direct detection of RNA at a fM sensitivity level without a target RNA amplification indicates that the Cas13a system is one of the most sensitive detection assays currently known. Furthermore, we introduced the transcription process before the CRISPR/Cas13a assay, using T7 promoter tagged DNA instead of RNA to avoid any instability problem during the procedure of incubation and washing. CRISPR/Cas13a was applied to detect the DNA transcripts for further enhancement of the sensitivity. As shown in Figure 1E, the T7 transcription process was added prior to the CRISPR/Cas13a assay. The results show that CRISPR/Cas13a is capable of detecting DNA transcripts at concentrations as low as 500 aM (as shown in Figure 1F, G), and Figure 1G presents an enlarged view of the low concentration curve in Figure 1F. The LOD of post-transcriptional detection was enhanced by two orders of magnitude compared to direct RNA detection.

**Figure 1.**
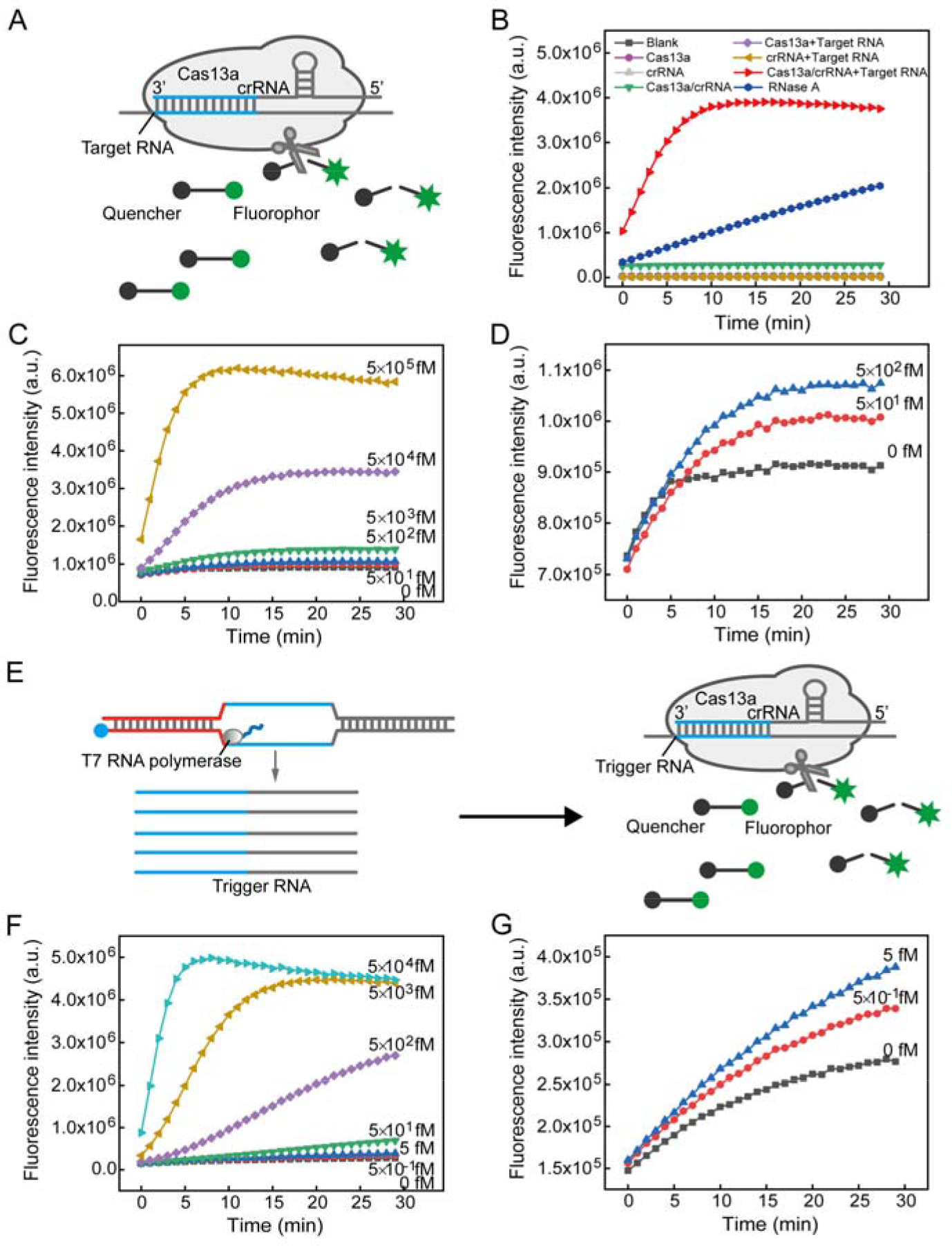
(A) Schematic for the principle of a Cas13a/crRNA-mediated RNA triggered signal amplification system. (B) Fluorescence measurement of LbuCas13a activity. RNase A was used as a positive control for the degradation of the RNA reporter probe. (C) Sensitivity of Cas13a/crRNA-mediated target RNA detection. Real-time fluorescence kinetic measurement of Cas13a reactions initiated by target RNA concentrations from 50 to 5×105 fM. (D) An enlarged view of the curves at the low concentrations of 0, 50, and 500 fM in Figure C. Data represent mean ± s.d., n = 3, three technical replicates. (E) Schematic for the principle of a Cas13a/crRNA-mediated RNA triggered signal amplification system after DNA transcription. (F) Sensitivity of Cas13a/crRNA-mediated RNA detection after DNA transcription. Real-time fluorescence kinetic measurement of Cas13a reactions initiated by transcription of DNA concentrations from 0.5 to 5×10^4^ fM. (G) An enlarged view of the curves at the low concentrations of 0, 0.5, and 5 fM in Figure F. Data represent mean ± s.d., n = 3, three technical replicates.

The achieved impressive sensitivity enabled us to construct a new ELISA built on the basis of a transcription assisted CRISPR/Cas13a assay. As is well known, classical ELISA is a heterogeneous assay format using a solid phase well plate. We next proved the feasibility of utilizing DNA transcription for this purpose by using a 96-well plate. As shown in Figure 2A, streptavidin was used to directly coat the 96-well plates, and then the plates were blocked with 1% BSA protein. A biotinylated DNA amplification template containing a T7 promoter sequence at one end was then added to the plate. Then, the unbound DNA amplification template was removed by washing. Next, transcription reaction buffer, T7 RNA polymerase, and nucleotide triphosphates (NTPs) were mixed together and transcribed at 37 °C for 1 h. Finally, the transcription products were detected by CRISPR/Cas13a. The fluorescence kinetic curves of each well were recorded, and the fluorescence intensity increased at 50 pM of template DNA (Figure 2B, red curve). In addition, the results were also expressed by the calibration values (*Δ*τ) in Figure 2C. In the presence of 50 pM template DNA the *Δ*τ value was much stronger than that of the negative control, indicating that the template DNA was successfully ligated to the plate and the DNA transcript can be successfully detected in a solid phase format.

**Figure 2.**
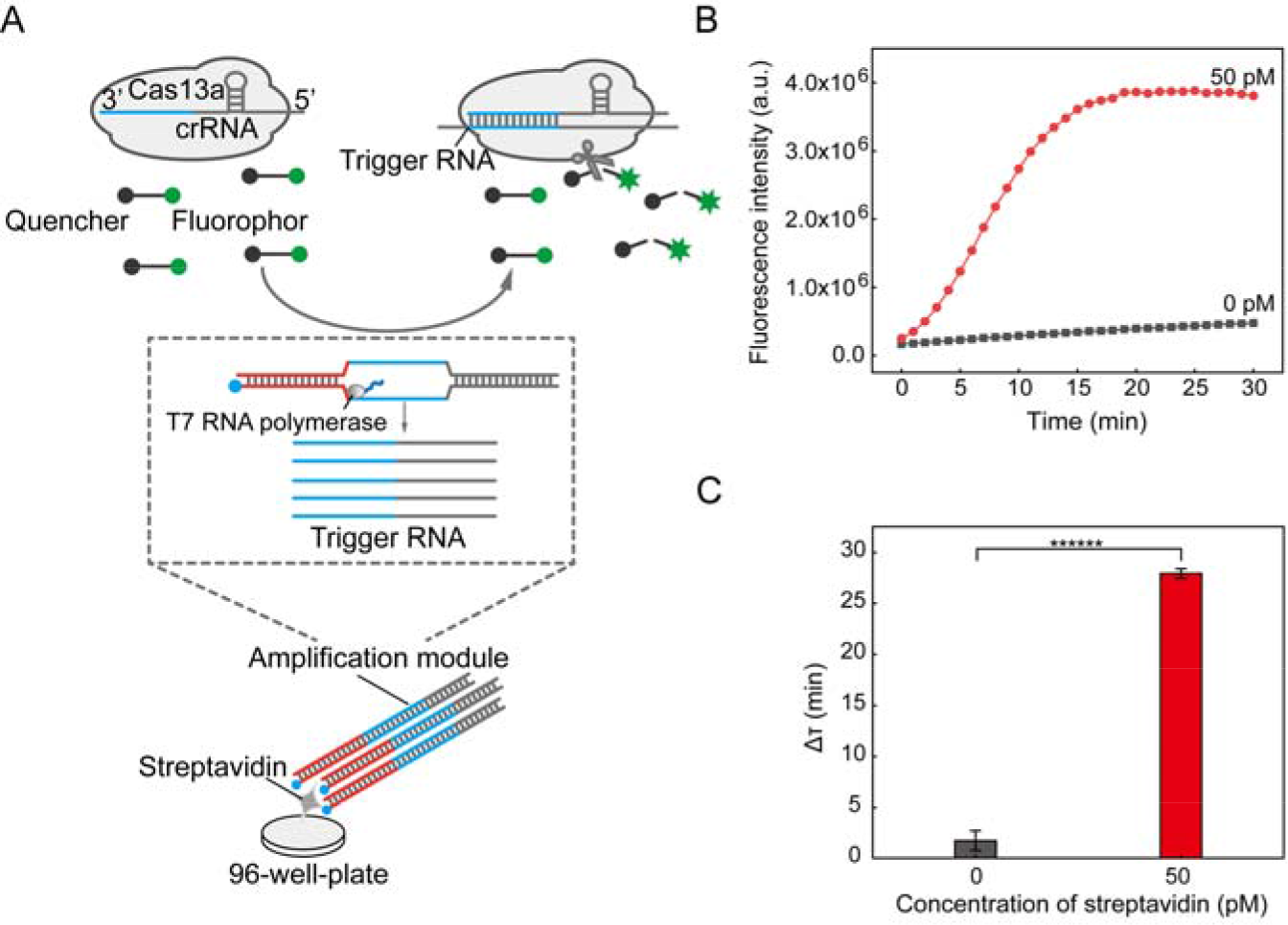
Validation of the compatibility of the Cas13a/crRNA-mediated RNA detection system with solid phase DNA transcription. (A) Streptavidin was precoated on a 96-well plate. The biotin-dsDNA amplification template (the amplification module) then bound to the streptavidin. The bound biotin-dsDNA was used as the template for DNA transcription by T7 RNA polymerase. (B) Real-time fluorescence kinetic measurement of simplified CLISA. The threshold was set to determine the critical time τ, which is the minimal time to reach the threshold. A calibration curve was then established by plotting *Δ*τ(*Δ*τ = 30 min − τ) as a function of the concentrations of antigen (C) (Student’s t-test; ******P < 0.00001). Data represent mean ± s.d., n = 3, three technical replicates.

After demonstrating a solid phase transcription assisted CRISPR/Cas13a assay, CLISA was developed by utilizing streptavidin as a bridge to link biotinylated detection antibodies to biotinylated DNA amplification templates, followed by DNA transcription to produce trigger RNA. The Cas13a system was then employed to detect the product, and when a trigger RNA is present, a substantial number of signal probes can be cleaved for a second signal amplification. In the CLISA, antigen-antibody binding, template transcription and Cas13a detection were all performed at 37 °C.

We chose human IL-6 and human VEGF as models to validate the CLISA. Human IL-6 is an inflammatory factor produced by tumor cells, T cells, and lymphocytes^25–26^. Human VEGF is involved in the pathogenesis and progression of many angiogenesis-dependent diseases, including cancer, certain inflammatory diseases, and diabetic retinopathy^27^. Human IL-6 and human VEGF have been considered to be important factors in disease development.

First, we applied CLISA to detect human IL-6. Serially diluted human IL-6 antigen and biotinylated detection antibody were added sequentially to form 'antibody-antigen-antibody' complexes. After that, streptavidin and the biotinylated DNA amplification template, which has been optimized as shown in Figure S3, were added sequentially, resulting in binding of the DNA amplification template to the 'antibody-antigen-antibody’ complex. Unbound templates were washed away, and then T7 RNA polymerase was utilized to amplify the amplification template (Figure 3A). As shown in Figure 3B, it is noted that the *Δ*τ is linear with the logarithm of human IL-6 concentrations in the range from 160 fg/ mL (8 fM) to 0.1 ng/mL (5 pM), and the linear regression equation is *Δ*τ = 8.496 lg *C* − 14.112 (R^2^ = 0.989) with a LOD of 45.81 fg/mL(2.29 fM). In addition, a commercial human IL-6 ELISA kit was subjected to the same experiment and showed a LOD of 12.09 pg/mL (605 fM) (blue curve). It is significant that the sensitivity of CLISA was 264-fold higher than that of the commercial ELISA kit.

**Figure 3.**
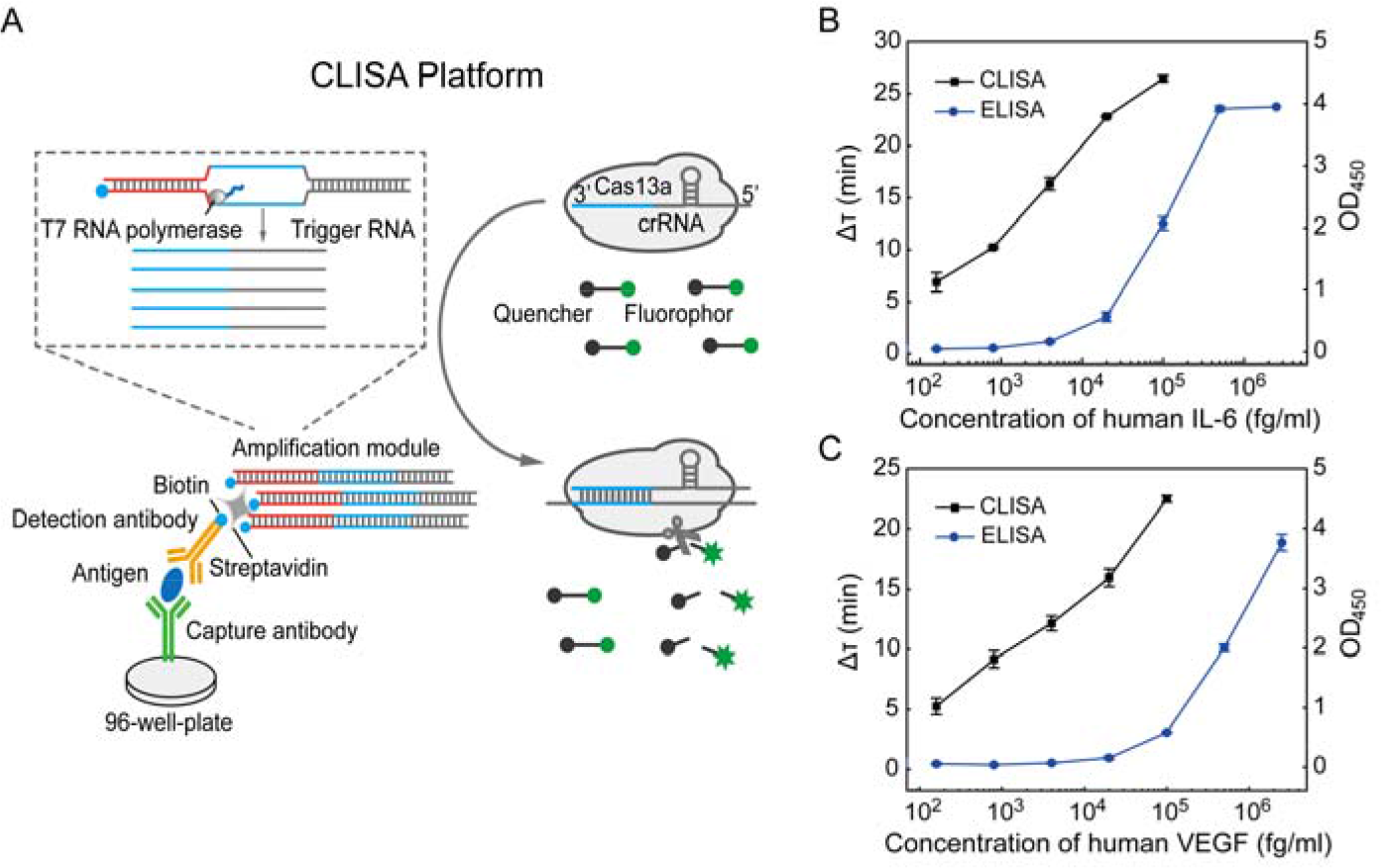
(A) Schematic for the principle of CLISA. In a CLISA assay, the capture antibody first binds to the antigen of interest. A detection antibody, which binds to a distant, nonoverlapping epitope in the antigen, is biotinylated and linked to a biotin-dsDNA template (the amplification module) through streptavidin. T7 RNA polymerase is then used to amplify the DNA template, producing many copies of RNA substrate, the amount of which is representative of the original amount of antigen. (B) Detection of human IL-6. Human IL-6 was added to the coated plate at a series of five-fold dilutions from 160 fg/mL (8 fM) to 100 pg/mL (5 pM). A parallel ELISA experiment was also performed with a series of fivefold dilutions from 160 fg/mL (8 fM) to 2.5 ng/mL (125 pM). Data represent mean ± s.d., n = 3, three technical replicates. (C) Detection of human VEGF. Human VEGF was added to the coated plate at a series of five-fold dilutions from 160 fg/mL (4 fM) to 100 pg/mL (2.5 pM). A parallel ELISA experiment was also performed at a series of fivefold dilutions from 160 fg/mL (4 fM) to 2.5 ng/mL (62.5 pM). Data represent mean ± s.d., n = 3, three technical replicates.

In addition, as displayed in Figure 3C, we also applied CLISA to detect human VEGF. In the range of 160 fg/mL (4 fM) to 0.1 ng/mL (2.5 pM) of human VEGF, there is a linear relationship between the *Δ*τ and the logarithm of human VEGF concentrations, with a linear regression equation of *Δ*τ = 6.347 lg *C* − 9.577 (R^2^ = 0.985) and a LOD as low as 32.27 fg/mL (0.81 fM). The commercial ELISA human VEGF kit showed a LOD of 20 pg/mL (500 fM) (Figure 3C). This result indicated that the LOD of the CLISA was also reduced by 617-fold compared to the commercial ELISA kit.

Next, we evaluated the analytical potential of the CLISA method for complex samples (Table 1). We added human IL-6 to diluted mouse serum (20%) to demonstrate whether the CLISA method is as resistant to matrix interference as a conventional ELISA method. The recovery test evaluating from three concentrations (100, 20, and 4 pg/mL) of human IL-6 samples showed that the recoveries were 96.21%, 101.32%, and 104.15%, respectively. Since the procedure of the current CLISA method requires washing similar to the conventional ELISA method, it is not surprising that the CLISA method has achieved an excellent anti-interference ability.

**Table 1.**
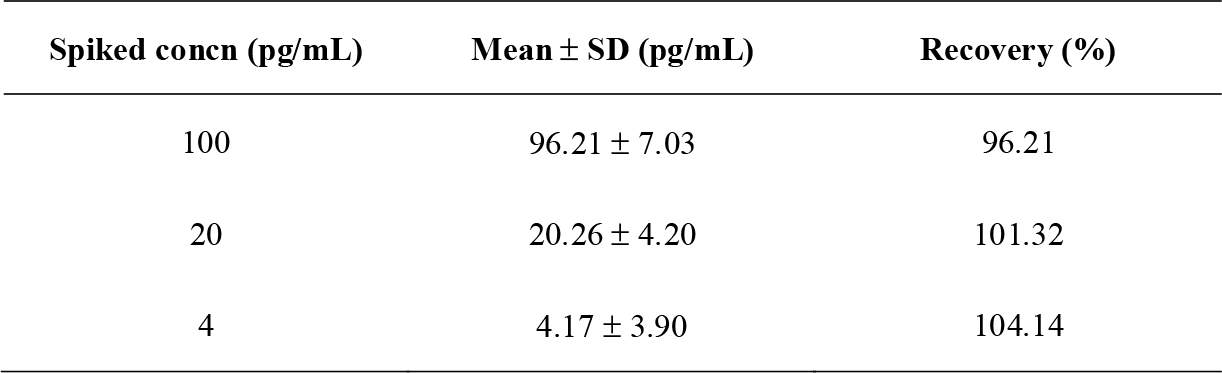
Recovery experiments of human IL-6 in serum samples

| Spiked concn (pg/mL) | Mean ± SD (pg/mL) | Recovery (%) |
| --- | --- | --- |
| 100 | 96.21 ± 7.03 | 96.21 |
| 20 | 20.26 ± 4.20 | 101.32 |
| 4 | 4.17 ± 3.90 | 104.14 |

Compared with the traditional ELISA, CLISA adds a transcription process to expand the target, and the signal is further enhanced by the collateral cleavage activity of CRISPR/Cas13a. As a result, the sensitivity of CLISA can be effectively ameliorated by two-step amplification. Furthermore, the whole process of the CLISA is performed at 37 °C, which is an isothermal process without the need for a thermal cycling program. It is worth noting that the CLISA procedure is completely compatible with existing commercial ELISA equipment. Although the experiments herein were performed manually, it is obvious that this method is compatible with current high throughput liquid handling robots for washing plates and reagent dispersion. Due to its improved sensitivity over commercial ELISA kits and its adaptability to high throughput and automation technologies, CLISA is able to detect low-abundance proteins that conventional ELISA cannot. In addition, we compared CLISA with several other immunological methods (Table 2). This work shows that our CLISA method is superior in sensitivity to most of the reported amplification strategies. Although the T7 transcription amplification assay reports a better sensitivity, the CLISA method demonstrates superior linearity and speed.

**Table 2.** Comparison of protein test results across published reports.

| Analytical methods | Sensitivity | Dynamic<br>range | Isothermal | Time to<br>result | Additional<br>instrument | Target |
| --- | --- | --- | --- | --- | --- | --- |
| CLISA (this work) | 0.8 fM | 3-logs | √ | 4.5 h | × | IL-6,<br>VEGF |
| ELISA (this work) | 12 pM | 3-logs | √ | 3.5 h | × | IL-6,<br>VEGF |
| Nanozyme<sup>7</sup> | 835 fM | 2-logs | √ | 1.4 h | √ | CEA |
| Nanoprobe<sup>9</sup> | 33 fM | 3-logs | √ | 2 h | × | IL-6 |
| Immune-PCR<sup>12</sup> | 5 fM | 5-logs | × | 5 h | √ | VEGF |
| Immune-HCR<sup>16</sup> | 500 fM | 3-logs | × | 3 h | √ | IL-2,9,10 |
| Immune-RCA<sup>14</sup> | 50 fM | 3-logs | × | 3 h | √ | IL-6 |
| Proximity ligation<br>assay<sup>18</sup> | 10 fM | 5-logs | × | 2 h | √ | VEGF |
| T7 transcription<br>amplification<sup>19</sup> | 0.08 fM | 3-logs | √ | 7 h | × | Her2 |

## CONCLUSIONS

In summary, we developed a highly sensitive, isothermal method for detecting low-abundance proteins based on the collateral cleavage activity of CRISPR/Cas13a initiated by trigger RNA. The sensitivity of CLISA was effectively improved by the amplification of T7 transcription and the collateral cleavage activity of CRISPR/Cas13a. Using human IL-6 and human VEGF as model analytes, the sensitivity of CLISA has been drastically boosted, with a LOD as low as 45.81 fg/mL (2.29 fM, 264-fold improvement) and 32.27 fg/mL (0.81 fM, 617-fold improvement) compared to commercialized ELISA kits. Moreover, the method is a compatible, automated and high-throughput assay that allows for rapid screening of large numbers of samples simultaneously, providing potential ultrasensitive detection methods for biosensing, medical research, and molecular diagnostics.

## ASSOCIATED CONTENT

Supporting Information

## ACKNOWLEDGMENTS

We thank Professor Yanli Wang for supplying the plasmid for LbuCas13a expression. This work was supported by the National Natural Science Foundation of China (Grant 21475048; 21874049, 21904042, 21675067), the National Science Fund for Distinguished Young Scholars of Guangdong Province (Grant 2014A030306008), the National Key Research and Development Program of China (Grant 2016YFD0501300), and the Natural Science Foundation of Jiangsu Province (No. BE2019645), and the Foundation of Postgraduate Research and the Practical Innovation Program of Jiangsu Normal University (No. 2018YXJ124).

