## Supporting information for "CRISPR/Cas13a signal amplification linked immunosorbent assay (CLISA)"

**1. Supplementary Table**

**Table S1.** Sequences used in this study

| **Name** | **Sequences (5’ to 3’)** |
| --- | --- |
| DNA1 | GAAATTAATACGACTCACTATAGGG CGCCTGAACCACCAGGCTATATCTGCCACTTT |
| DNA2 | Biotin-AAAGTGGCAGATATAGCCTGGTGGTTCAGGCGCCCTATAGTGAGTCGTATTAATTTC |
| crDNA1 | GAAATTAATACGACTCACTATAGGGGGCCACCCCAAAAATGAAGGGGACTAAAACACAGATATAGCCTGGTGGTTC |
| crDNA2 | GAACCACCAGGCTATATCTGTGTTTTAGTCCCCTTCATTTTTGGGGTGGCCCCCTATAGTGAGTCGTATTAATTTC |

**2. Supplementary Figures**

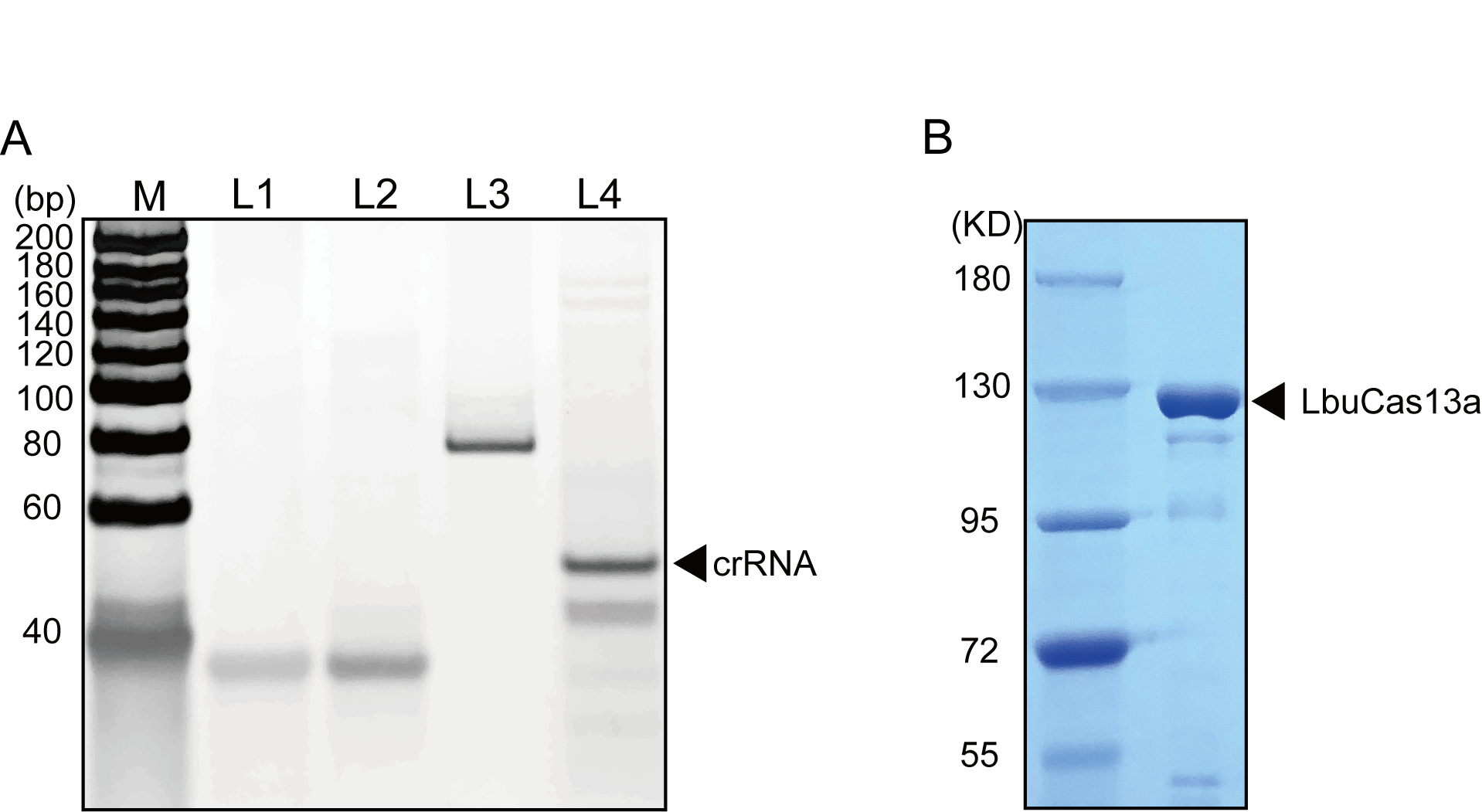

**Figure S1.** (A) PAGE gels of crRNA. M: DNA marker, L1: crDNA1; L2: crDNA2; L3: ds-crDNA; L4: crRNA. (B) SDS-PAGE gels of purified LbuCas13a.

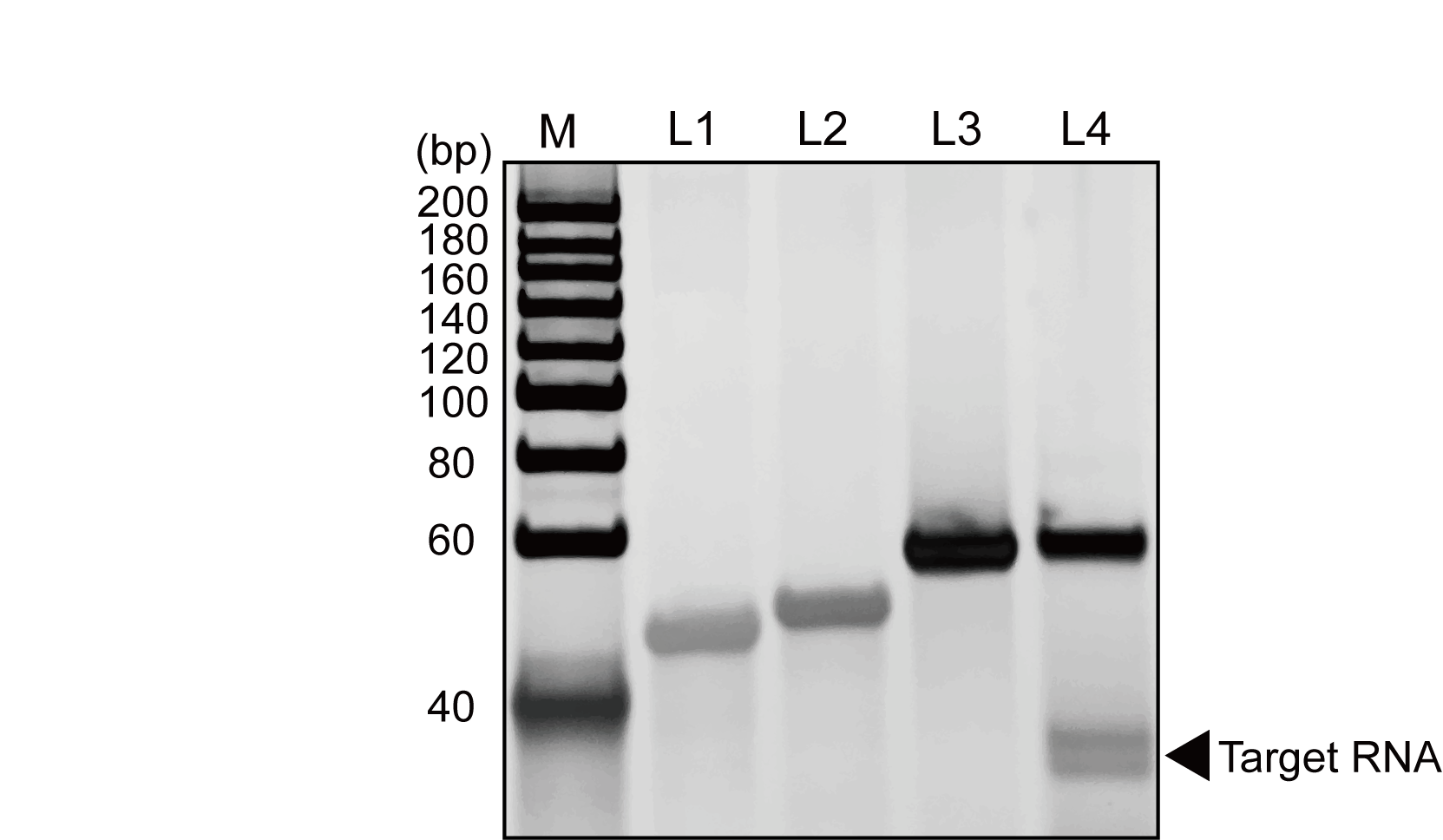

**Figure S2.** PAGE gels of target RNA. M: DNA marker, L1: DNA1; L2: DNA2; L3: ds-DNA; L4: Target RNA.

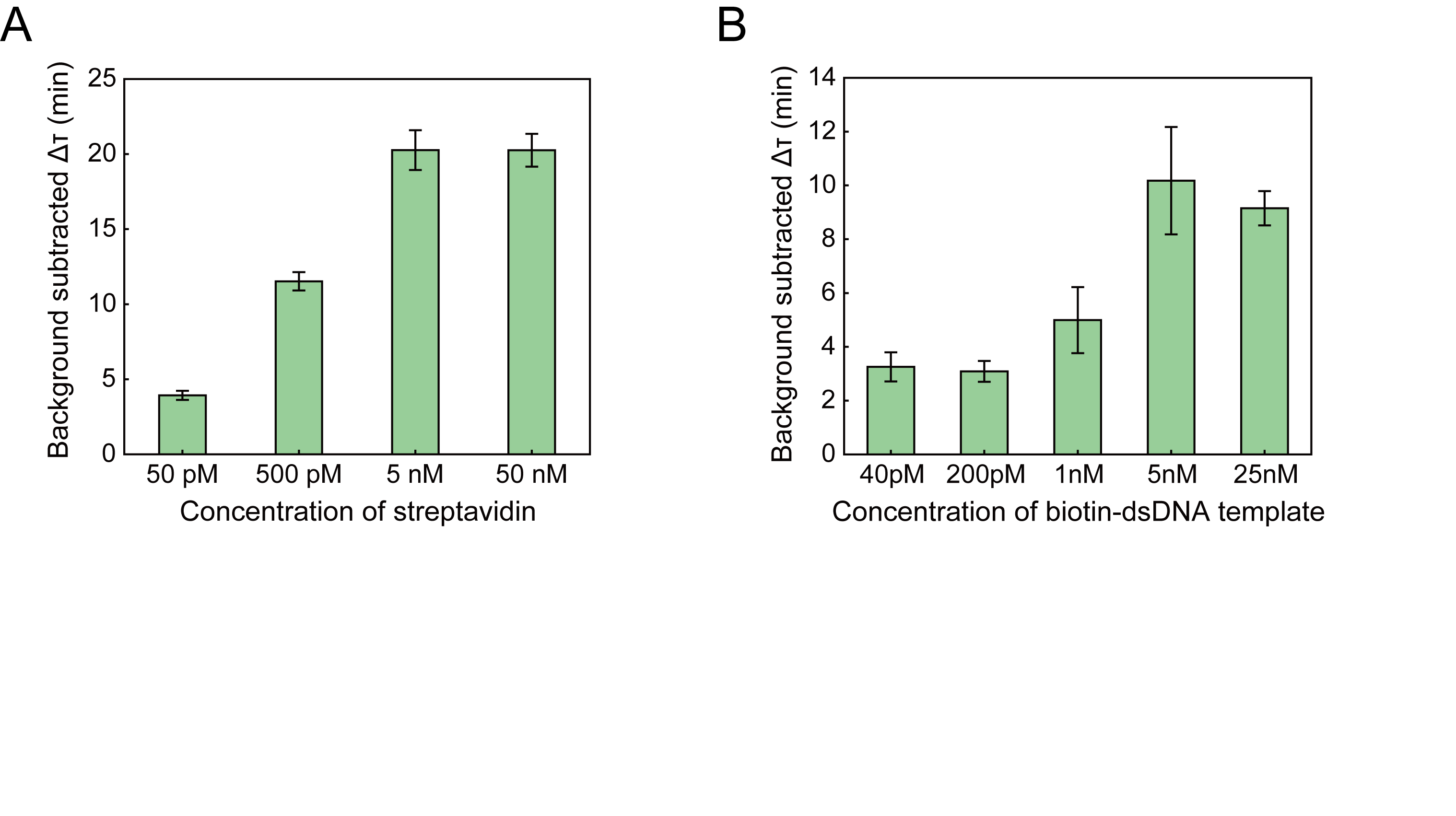

**Figure S3.** (A) Optimization of the streptavidin concentration from 50 pM to 50 nM. The optimal concentration is 5 nM. (B) Optimization of the biotin-dsDNA template concentration from 40 pM to 25 nM. The optimal concentration is 5 nM. Data represent mean ± s.d., n = 3, three technical replicates.
